# Functional metagenomics reveals novel β-galactosidases not predictable from gene sequences

**DOI:** 10.1101/047167

**Authors:** Jiujun Cheng, Tatyana Romantsov, Katja Engel, Andrew C. Doxey, David R. Rose, Josh D. Neufeld, Trevor Charles

**Affiliations:** Department of Biology, University of Waterloo, 200 University Avenue West, Waterloo, ON, N2L 3G1, Canada

**Keywords:** Functional metagenomics, soil metagenomic library, β-galactosidase, Sinorhizobium meliloti

## Abstract

A soil metagenomic library carried in pJC8 (an IncP cosmid) was used for functional complementation for β-galactosidase activity in both *α-Proteobacteria (Sinorhizobium meliloti)* and *γ-Proteobacteria (Escherichia coli).* One *β*-galactosidase, encoded by overlapping clones selected in both hosts, was identified as a member of glycoside hydrolase family 2. ORFs obviously encoding possible β-galactosidases were not identified in 19 other clones that were only able to complement *S. meliloti.* Based on low sequence similarity to known glycoside hydrolases but not β-galactosidases, three ORFs were examined further. Biochemical analysis confirmed that all encoded *β*-galactosidase activity. Bioinformatic and structural modeling implied that Lac161_ORF10 protein represented a novel enzyme family with a five-bladed propeller glycoside hydrolase domain.

## Introduction

Soils harbour the greatest genetic diversity of any habitats on Earth (Curtis et al. 2002). Our knowledge of microorganisms comprising soil communities is hampered by cultivation challenges for many microorganisms in these communities (Simon and Daniel 2011), although improvements in cultivation methods are addressing this bottleneck (Shade et al. 2012). The genomes of metabolically versatile soil microbes are potential sources of biocatalysts for use in various industrial processes. Limited knowledge of links between sequence and function prevent rapid progress in bioinformatics-based systems biology. As a result, metagenomics can be used to explore the collective genetic constituency of environmental microbes, including those that are difficult to culture through conventional microbiological techniques. Sequence-based and function-based strategies are used in metagenomics, depending on the main objectives of the particular study. Sequence-based metagenomics identifies genes by sequence similarity to known database sequences. However, it is difficult, if not impossible, to reliably predict the function of truly novel genes without experimental evidence. Functional screening strategies are based on phenotypic detection of the desired activity, heterologous complementation of host strains, and induced gene expression (André et al. 2014) (Simon and Daniel 2011; Taupp et al. 2011). These experimental activities have identified novel genes showing little similarity to genes of known function (Beloqui et al. 2010; Ferrer et al. 2009; Iqbal et al. 2012) (Ufarté et al. 2015). In addition, heterologous complementation screening strategies facilitate simultaneous screening of millions of metagenomic clones. Most functional screens are performed in *Escherichia coli* of the *γ-Proteobacteria*. Because gene expression is often host-dependent (Gabor et al. 2004), multi-host systems increase the likelihood of the successful gene expression (Aakvik et al. 2009; Craig et al. 2010; Hao et al. 2010; Martinez et al. 2004; Taupp et al. 2011; Wang et al. 2006) (Cheng et al. 2014; Li et al. 2005; Ly et al. 2011) (Biver et al. 2013).

Glycoside hydrolases (GH) hydrolyze the glycosidic linkages of glycosides and oligosaccharides, and are classified into 131 families based on the similarity of amino acid sequences (Cantarel et al. 2009); www.cazy.org/Glycoside-Hydrolases.html). The β-galactosidase (EC 3.2.1.23) enzymes are grouped within GH1, GH2, GH35, GH42 and GH59 families. β-Galactosidase hydrolytic activity is used in applications such as reducing the lactose content in dairy products (Harju et al. 2012), producing bioethanol from cheese whey (Guimaraes et al. 2010), and detecting lactose as a biosensor (Marrakchi et al. 2008). The transgalactosylation activity is used to synthesize galactosylated products (Gosling et al. 2010). Functional screening of metagenomic libraries resulted in discovery of a GH43 enzyme acting on multiple substrates including lactose (Ferrer et al. 2012), cold-adapted or thermostable GH42 β-galactosidases (Wang et al. 2010; Zhang et al. 2013), a thermostable-alkalophilic or cold-active GH1 (Gupta et al. 2012; Wierzbicka-Wos et al. 2013), and two novel β-galactosidases without any similarity to known GHs (Beloqui et al. 2010).

In this study, we demonstrate the value of metagenomic cosmid libraries for enzyme discovery. Using lactose as the sole carbon source to support growth of *Sinorhizobium meliloti*, we identified three new β-galactosidases from one of the soil libraries, and characterized the biochemical properties of these novel enzymes. These new enzymes represent new associations of protein sequence space with this substrate specificity.

## Materials and Methods

### Bacterial strains, plasmids, cosmids, and growth conditions

Several bacterial strains, plasmids, and cosmids were used in this study (Table 1). All *E. coli* strains were grown at 37°C in LB medium (1% tryptone, 0.5% yeast extract and 0.5% NaCl, pH.). *S. meliloti* strains were grown at 30°C in LB supplemented with 2.5 mM CaCl_2_ and 2.5 mM MgSO_4_ (LBmc; (Finan et al. 1984)). Antibiotics were used at the following final concentrations: streptomycin (100 (μg/ml for *E. coli*, 200 (μg/ml for *S. meliloti)*, neomycin (200 (μg/ml), rifampicin (100 (μg/ml), kanamycin (50 (μg/ml), tetracycline (20 (μg/ml for *E. coli*, 10 (μg/ml for *S. meliloti)*, gentamicin (10 (μg/ml).

**Table 1.** Bacterial strains, plasmids, and cosmids.

| Bacteria, plasmids, cosmids | Characteristics | References |
| --- | --- | --- |
| **Bacteria** |  |  |
| ***S. meliloti*** |  |  |
| Rm1021 | SU47 *str-21*, Sm<sup>R</sup> | (Meade et al. 1982) |
| RmF728 | Rm1021 derivative (*lacEFGZ1K1*), Sm<sup>R</sup> Nm<sup>R</sup> | (Charles and Finan 1991) |
| SmUW253 | Rm1021 derivative (*lacZ1*), Sm<sup>R</sup> | This work |
| ***E. coli*** |  |  |
| DH5α | F<sup>-</sup> φ80*lacZ*ΔM15 Δ(*lacZYA-argF*) U169 *recA1 endA1 hsdR17 phoA supE44 thi-1 gyrA96 relA1* | (Hanahan 1983) |
| DH5α (Rif<sup>R</sup>) | A spontaneous Rif<sup>R</sup> mutant of DH5α, Rif<sup>R</sup> | (Cheng et al. 2014) |
| HB101 | F<sup>-</sup> *supE44 lacY1 ara-14 galK2 xyl-5 mtl-1 leuB6 recA13 rpsL20 thi-1 proA2 hsdSB20*, Sm<sup>R</sup> | (Boyer and Roulland-Dussoix 1969) |
| BL21(DE3)pLysS | F<sup>-</sup> *ompT lon hsdS<sub>B</sub> gal dcm λ*(DE3) pLysS, Cm<sup>R</sup> | Novagen |
| MT616 | *pro82 thi-1 endA hsdR17 supE44 recA56* (pRK600), Cm<sup>R</sup> | (Finan et al. 1984) |
| **Plasmids and cosmids** |  |  |
| pET-30a(+) | Expression vector, Km<sup>R</sup> | Novagen |
| pET-30b(+) | Expression vector, Km<sup>R</sup> | Novagen |
| pK19mobsacB | Cloning vector, Km<sup>R</sup> | (Schäfer et al. 1994) |
| pSRKGm | pBBR1MCS-5 derivative, Gm<sup>R</sup> | (Khan et al. 2008) |
| pRK600 | Conjugation helper plasmid, Cm<sup>R</sup> | (Finan et al. 1984) |
| pJC8 | Low-copy broad-host-range Gateway<sup>®</sup> entry cosmid, Tc<sup>R</sup> Gm<sup>R</sup> | (Cheng et al. 2014) |
| pJC44 | pK19mobsacB carrying in-frame deletion of Rm1021 *lacZ1*, Km<sup>R</sup> | This work |
| pJC97 | pJC98 carrying Lac36W-ORF11, Gm<sup>R</sup> | This work |
| pJC98 | pSRKGm derivative carrying a His-tag region from pET-30b(+), Gm<sup>R</sup> | This work |
| pJC102 | pJC98 carrying Lac161-ORF7, Gm<sup>R</sup> | This work |
| pTR5 | pET-30a(+) carrying Lac161-ORF10, Km<sup>R</sup> | This work |

### Functional screening of β-galactosidases

Cosmids carrying metagenomic DNA of corn field soil (12AC; (Cheng et al. 2014)) were isolated from the pooled library clones using GeneJET Plasmid Miniprep Kit (Thermo Scientific). *E. coli* DH5α *(lacZYA)* was grown in LB to an OD_600_ of 0.6. Cells were collected by centrifugation at 4°C and at 12,300 x *g* for 20 min, washed three times with cold 10% glycerol (equal vol, ½ vol and 1/10 vol, respectively). Cells were gently suspended in 2 ml of ice-cold 10% glycerol (about 3 x 10 cells/ml). Electrocompetent cell volumes of 40 μl were mixed with 1 μl of cosmid DNA (45 ng) in a cold 1.5-ml microtube on ice, then transferred to a cold electroporation cuvette (0.2 cm, Bio-Rad). Electroporation was performed using Gene Pulser (Bio-Rad; C = 25 uF; PC = 200 ohm; V=3.0 kV). Liquid SOC medium (1 ml) was added to the cuvette after one pulse. Electroporated cells were transferred to a 1.5-ml microcentrifuge tube and incubated at 37°C in a water bath for 30 min, inverting the tube every 5 min. The tube was then shaken at 3 C and at 200 rpm for 30 min. Following concentration by centrifugation, cells were spread on LB Tc plates, and incubated overnight at 37°C. Multiple electroporations were performed to obtain the desired numbers of recombinant *E. coli* DH5α clones. Transformants were pooled and saved at −75°C after addition of DMSO (7% final concentration).

The pooled *E. coli* DH5α cosmid clones were washed three times with 0.85% NaCl and then spread on defined M9 medium (Cheng et al. 2007) supplemented with L-arginine (50 μg/ml), thiamine (10 μg/ml), tetracycline (15 μg/ml), with lactose (15 mM) as the sole carbon source, as well as the chromogenic substrate X-gal (36 μg/ml). Plates were incubated at 37°C for 1-3 days. Positive blue colonies were streak purified once on M9-lactose plates. The Lac^+^ clones were inoculated in 3 ml of LB Tc medium and grown overnight at 37°C. Cosmid DNA was isolated using the GeneJET Plasmid Miniprep Kit (Thermo Scientific), digested simultaneously with EcoRI-BamHI-HindIII, then resolved on 1% agarose gels. Cosmids representative of distinct restriction patterns were retransformed into *E. coli* DH5α, and then spread on the M9-lactose to confirm the Lac^+^ phenotype.

To screen for Lac^+^clones in *S. meliloti*, 12AC cosmids were conjugated from *E. coli* DH5α into *S. meliloti* RmF728 *(lacEFGZ1K1;* (Charles and Finan 1991)) with helper plasmid pRK600. The pooled 12AC library clones of 0.25 ml were mixed with 2 ml each of overnight-grown *S. meliloti* RmF728 and *E. coli* DH5α (pRK600). Cells were collected by centrifugation at 12,300 x *g* for 3 min, washed twice with 2 ml of 0.85% NaCl, then resuspended in 0.5 ml of 0.85% NaCl. Mixed cells were spotted on LB agar and incubated overnight at 30°C. Following collection of the mating spot in a 2.0-ml microtube and washing twice with 0.85% NaCl, the conjugation mixture was serially diluted and plated on the defined M9 medium (Nm Tc) supplemented with biotin (0.3 (μg/ml), thiamine (10 (μg/ml), X-gal (36 (μg/ml), and lactose (15 mM) as the sole carbon source. Lac^+^ colonies were streak purified once on M9 lactose plates. Cosmids were then transferred from *S. meliloti* to *E. coli* DH5α (Rif^R^) via conjugation with the helper plasmid pRK600. *E. coli* DH5α carrying the empty cosmid pJC8 was used as a negative control. Lac^+^ cosmid DNA was prepared and analyzed by EcoR-BamH-HindIII digestion as described previously.

To further verify the Lac^+^ phenotype conferred by the complementing clones, we constructed a *S. meliloti* strain SmUW253 in which the *lacZ1* gene encoding a β-galactosidase was deleted but an ABC-type transporter of lactose encoded by *lacEFGK1* is functional. A 5’- fragment upstream of the *lacZ1* gene was PCR amplified using the genomic DNA of *S. meliloti* Rm1021 as a template and primers JC98 and JC99 (Table S1). A 3’-region downstream of the *lacZ1* was obtained by PCR amplification using the same template and primers JC100 and JC101. The two PCR products of correct sizes were purified on an agarose gel, and then mixed in equal amount to serve as templates for the second PCR using primers JC98 and JC101 (Table S1). The precise deletion of the *lacZ1* ORF was then inserted into the EcoRI and HindIII sites in pK19mobsacB, yielding plasmid pJC44. Single cross-over recombination of pJC44 into *S. meliloti* Rm1021 genome was performed via conjugation. The double cross-over event (deletion of *lacZ1)* was selected by sucrose resistance, followed by screening for Km^S^ (loss of plasmid backbone) and white colonies on LB X-gal plate (deletion of LacZ1). Finally the *S. meliloti* Lac^+^ cosmids were conjugated from *E. coli* DH5α (Rif^R^) to the *S. meliloti* SmUW253 *(lac)* to confirm the Lac^+^ phenotype of the isolated 12AC cosmids.

The Lac^+^ phenotypes of *S. meliloti* strains were also verified by assaying β-galactosidase activity. Thirty-nine random Lac^+^ strains were grown overnight in LBmc, washed twice with 0.85% NaCl, and then subcultured (1:200 dilution) in M9 lactose medium. Following growth for 48 h, β-galactosidase activity was measured using o-nitrophenyl β-galactoside (ONPG) as described previously (Cowie et al. 2006).

### Cloning, expression, purification and characterization of β-galactosidases

The KOD Xtreme DNA polymerase (Novagen) was used for all PCR amplifications with several different primers (Table S1). PCR amplifications consisted of one cycle of 94°C for 5 min, 30 cycles of 94°C for 30 s, 50-57°C for 30 s, 68°C for 30 s to 3 min, and incubation at 68 C for 10 min. The Lac161_ORF10 was PCR amplified using primers lac161NdeI and lac161HindIII, and cloned into the NdeI-HindIII sites in pET-30a(+) to obtain pTR5. To clone putative GH genes with a C-terminal His tag in a broad-host-range plasmid, a 0.37-kb DNA fragment containing the NdeI site to the end of T7 terminator from pET-30b(+) was amplified using primers JC226 and JC227, and inserted into the NdeI-NheI sites in pSRKGm (Khan et al. 2008) to obtain plasmid pJC98. The Lac161_ORF7 was obtained by PCR amplification using primers JC220 and JC221, and then inserted into the NdeI-SalI sites in pJC98, yielding pJC102. Lac36W_ORF11 was PCR amplified using primer pair JC212 and JC213, and then cloned into the NdeI-XhoI sites in pJC98 to obtain plasmids pJC97. Plasmids were verified by restriction enzyme digestion analysis.

The expression plasmids pTR5, pJC97, and pJC102 were introduced into *E. coli* BL21(DE3)pLysS using the CaCl_2_ transformation method. Gene overexpression was induced by adding 0.1 mM IPTG at 20°C for 16 h. Cell pellets were resuspended in a lysis buffer (100 mM potassium phosphate; pH 7.4, 5 mM MgSO_4_, 30 μg/ml DNase, 1 mg/ml lysozyme, 2 mM β-mercaptoethanol, and 0.5 mM phenylmethylsulfonyl fluoride), incubated on ice for 30 min, and then disrupted by three passes through EmulsiFlex-C3 (Avestin Inc. Ottawa, Ontario) pressure cell at an internal cell pressure of 1.6 x 10^8^ Pa. His-tagged proteins were purified from supernatants of cell extracts under native conditions using Co^2+^-NTA affinity chromatography (Clontech Laboratories). Purified proteins were dialyzed twice at 4°C against 50 mM potassium phosphate and 10 mM Tris-HCl (pH 7.4).

Enzyme activities were measured using a Glucose Oxidase Activity Assay Kit (Sigma-Aldrich) for quantifying the amount of glucose produced upon the addition of enzyme at different substrate concentrations. Assays were carried out in 96-well microtiter plates containing substrate (0.5–15 mM), 100 mM MES buffer of pH 6.5 (Lac161_ORF10) or pH 6.0 (Lac161_ORF7 and Lac36W_ORF11) and enzyme (0-5 mM). Reactions were incubated at 37°C (Lac161_ORF10), 42°C (Lac36W_ ORF11), and 50°C (Lac161_ORF3) for 30 min and terminated with Tris-HCl (pH 7) to a final concentration of 1 M. Aliquots of glucose oxidase/peroxidase reagent (125 μl) were added to each well and left to develop at 37°C for 30 min. Absorbance was measured at 450 nm and compared with a standard glucose curve to determine the amount of glucose released. All reactions were performed in triplicate.

### Bioinformatic analysis

Illumina raw sequence data were assembled as described previously (Lam et al. 2014). Open reading frames were annotated using MetaGeneMark (Zhu et al. 2010). Functions of proteins were predicted by BLAST analysis against NCBI non-redundant protein sequences, Pfam (Finn et al. 2014), and CAZy analysis toolkit (Park et al. 2010). Transmembrane helices were predicted by the TMHMM Server v. 2.0 (www.cbs.dtu.dk/services/TMHMM). Signal peptide was predicted using SignalP 4.0 (Petersen et al. 2011). Conserved protein domains were searched against NCBI Conserved Domain Database and analyzed with CDTree (Marchler-Bauer et al. 2013). Protein structure was predicted with Phyre 2.0 (Kelley and Sternberg 2009). Taxonomic affiliations of cosmid inserts were assigned based on compositional classifier PhyloPythiaS (Patil et al. 2012).

### Protein homology search against metagenomic datasets

SSEARCH36 (Pearson and Lipman 1988) was used to search 158 metagenomes (32 aquatic, 76 human gut, 50 soil) for homologs to Lac161_ORF7, Lac161_ORF10, and Lac36W_ORF11, with an *E*-value threshold of 0.01. The database of metagenomes was compiled based on the set of aquatic and human gut metagenomes (Doxey et al. 2015) (Qin et al. 2010), as well as a variety of soil metagenomes obtained from the MG-RAST server (metagenomics.anl.gov/). Accession numbers for all datasets are available in Supplementary Table S4. For comparison, and to estimate a background level of protein abundance using a housekeeping gene, all metagenomes were also searched for metagenomic homologs of the *rpoB* protein using HMMer (hmmer.org/) as implemented in MetAnnotate (Petrenko et al. 2015). Metagenomes possessing fewer than 100 *rpoB* hits were discarded, as these datasets were too small to yield meaningful results.

## Results

### Functional screening of β-galactosidases

Cosmid clones expressing β-galactosidase genes were screened in metagenomic library 12AC (Cheng et al. 2014). Functional β-galactosidase enzymes hydrolyse lactose (galactose-β-1,4-glucose) to galactose and glucose, facilitating the growth of bacterial hosts (*lac*) on M9 minimal media when lactose is used as the sole carbon source (Cheng et al. 2014). Because both the library host *E. coli* HB101 (*lacY1*) and surrogate *S. meliloti* RmF728 (*lac*) are resistant to streptomycin, which would affect selection of transconjugants in *S. meliloti*, 12AC cosmids were transferred from *E. coli* HB101 (Sm^R^ Tc^R^) to DH5α *(lacZYA)* via electroporation. We obtained ∼8.2 x 10^5^ recombinant clones (Tc^R^) of *E. coli* DH5α, which was ∼10 fold greater than the number of original cosmid clones. A total of 161 blue colonies were recovered on the selection medium following spreading the *E. coli* DH5α clones on M9-lactose plate with X-gal. Mapping with an EcoRI-HindIII-BamHI restriction enzyme digest demonstrated that these 161 clones represented 17 different banding patterns.

We decided to employ *S. meliloti* from the *α-Proteobacteria* as a soil-dwelling surrogate host for screening in an effort to expand the range of recovered β-galactosidase-encoding clones. *S. meliloti* strain RmF728 is a derivative of the well studied Rm1021 that has been modified to carry a genomic deletion that removes the lactose metabolism genes (Charles and Finan 1991). The 12AC cosmids were transferred from *E. coli* DH5α to the *S. meliloti* RmF728 via *en masse* triparental conjugation, and 1052 Lac^+^ colonies that were recovered on M9-lactose medium demonstrated reliable growth after streak purification. The colony color of these clones on M9-lactose containing X-gal ranged from white to varying shades of blue. The measurement of β-galactosidase activities of 39 random *S. meliloti* clones grown in M9 lactose medium (Table S2) confirmed that the ability to grow on lactose as sole carbon source was due to cosmid clone-encoded β-galactosidase activity.

Each of the Lac^+^ clones was transferred from *S. meliloti* by triparental conjugation to *E. coli* DH5α (Rif^R^). Electrophoretic comparison of 291 randomly chosen cosmids digested with EcoRI-HindIII-BamHI demonstrated 208 distinct patterns (65%), which suggested that the use of *S. meliloti* as a surrogate host for this screen yielded a greater diversity of β-galactosidase genes than when *E. coli* was used. There was some overlap with the clones isolated by complementation of *E. coli* DH5α, with four restriction enzyme digestion patterns common to both screens. In general, the clones showing a Lac^+^ phenotype in both *E. coli* and *S. meliloti* exhibited higher activity (Table S2).

### Sequencing and annotation of Lac^+^ cosmids

We randomly chose 3 distinct *E. coli* and 22 distinct *S. meliloti* Lac^+^ cosmids for high-throughput sequencing (Table 2; (Lam et al. 2014)). The Lac100B, Lac112W, and Lac224 sequences were partially assembled, and no β-galactosidase was readily predicted from those sequences. Complete insert sequences and annotated ORFs of the other 22 cosmids have been deposited in GenBank (Table 2). Based on taxonomic analysis, the metagenomic DNA carried by these clones was predicted to originate from at least four different bacterial phyla (*Cytophaga, Thermomicrobia, Verrucomicrobia, α*-, *β*-, *γ*- and *δ-Proteobacteria).* Cloned metagenomic DNA was GC rich overall (53% to 71%, 64% average).

**Table 2.**
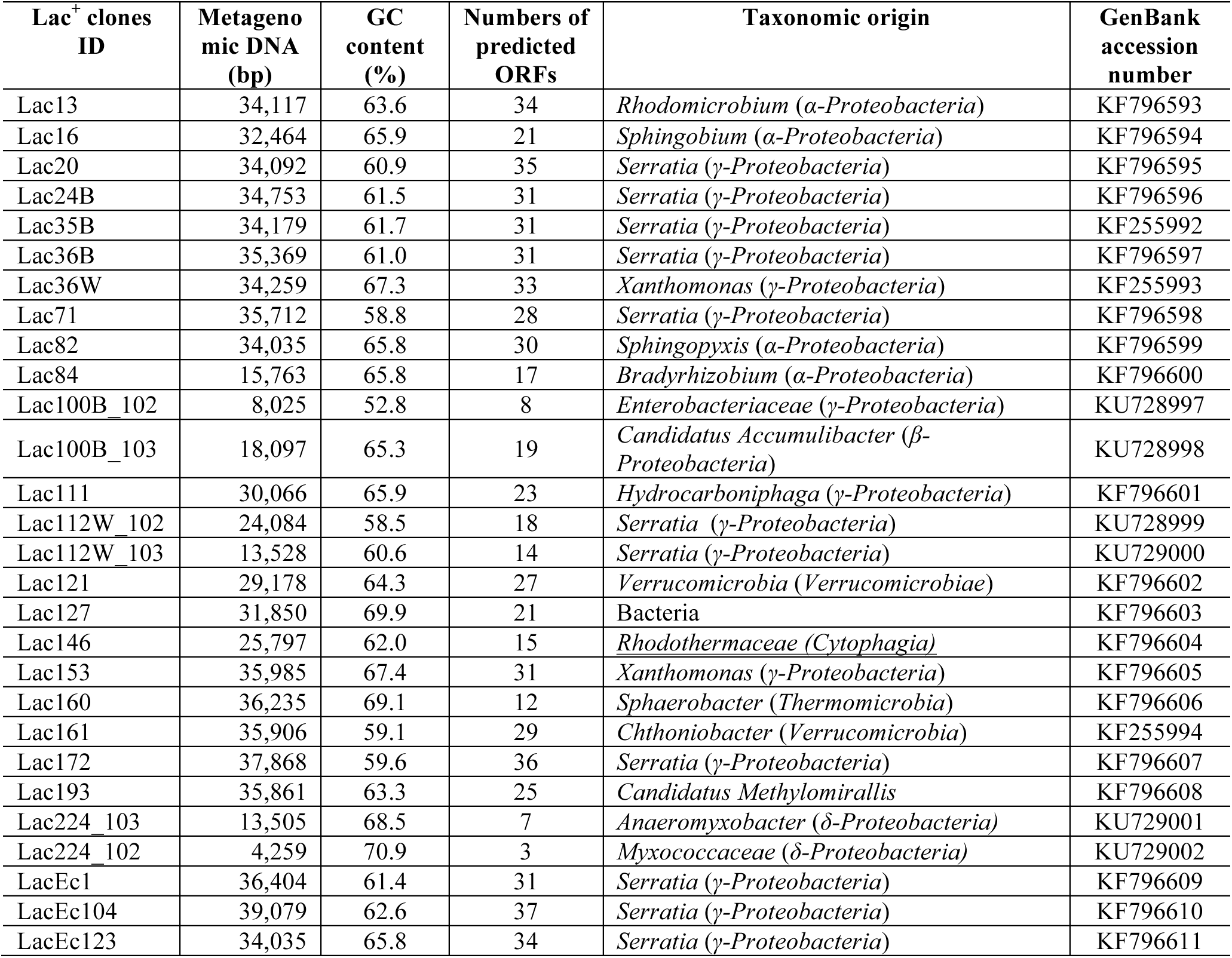
12AC metagenomic clones complementing *E. coli* DH5α (*lacZYA*) and *S. meliloti* RmF728 (*lacEFGZ1K1*) grown in M9-lactose medium. Cosmid sequences were obtained by Illumina sequencing. Taxonomic origin was determined using PhyloPythiaS.

The metagenomic DNA in *E. coli* Lac^+^ clones LacEc1, LacEc104, and LacEc123, and *S. meliloti* cosmid Lac24B, Lac36B, and Lac35B was predicted to originate from *Serratia* of the *γ-Proteobacteria* (Table 2). These clones overlapped over a segment of 15,344 bp (Fig. S1A). The 5’ region (positions 3 - 3,458) exhibited 93% identity to a chromosomal region (positions 2,604,251 - 2,607,707) of *Serratia marcescens* subsp. *marcescens* Db11 chromosome (GenBank accession HG326223), but the 3’ region (positions 6,652 - 15,344) of cloned DNA matched best to another region (positions 2,6143,370 - 2,623,056, 93% identity) of strain Db11. Eleven of 13 ORFs predicted in the overlapping region were 89-98% identical to the clustered orthologs (Fig. S1B). The second ORF (Lac35B, GenBank AGW45499) encodes a β-galactosidase (EC 3.2.1.23) with conserved domains of GH2 (Fig. 1; Fig. S1B). The enzyme matched to the predicted β-galactosidase (SMDB11_2462) of *S. marcescens* subsp. *marcescens* Db11 with 98% amino acid sequence identity. Additionally, the annotated β-galactosidase also shares 66% amino acid sequence identity to the well characterized β-galactosidase LacZ (GenBank, BAE76126) of *E. coli* K12 substr. W3110. The amino acid residues important for catalytic function in *E. coli* LacZ (Jacobson et al. 1994) are conserved in the annotated β-galactosidase at Glu^415^, His^417^, Glu^460^, Tyr^502^, and Glu^536^ (Fig. S2).

**Figure 1.**
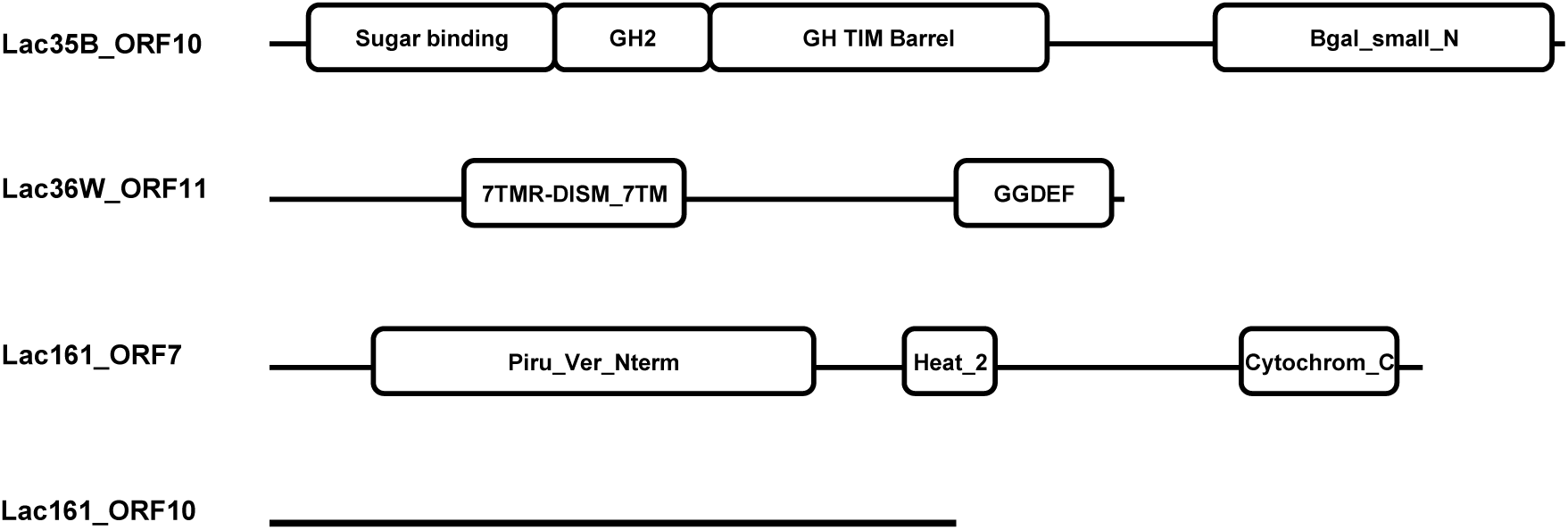
Conserved domains (*E*<0.01) in β-galactosidase isolated from 12AC metagenomic library clones.

Expression of the gene encoding the GH2 β-galactosidase from the cosmid clones in both *E. coli* and *S. meliloti* suggested a functional promoter(s) upstream of the gene. There were two regions homologous to the conserved −35 and −10 sites of RpoD promoters of *E. coli* (Lisser and Margalit 1993) and *S. meliloti* (MacLellan et al. 2006) (Fig. S1C) in the 102-bp intergenic region between the 3’ end (position 23) of an ORF (Lac35B_ORF9, GenBank AGW45500) encoding a two-component response regulator CitA, probably involved in Mg-citrate transport, and the β-galactosidase gene (Lac35B_ORF10, GenBank AGW45499) in Lac35B. These two putative promoters could drive expression of the β-galactosidase gene in *E. coli* and *S. meliloti*. Unlike the *E. coli lac* operon, there was no LacI homolog predicted in the cloned metagenomic DNA of those six cosmids. In addition, expression of the gene encoding β-galactosidase was neither inhibited by 15 mM glucose nor stimulated by addition of 0.4 mM IPTG in M9 medium.

Because the lactose permease LacY in *E. coli* DH5α (Meselson and Yuan 1968) and ABC-type transporter LacEFGK1 for lactose in *S. meliloti* RmF728 are deleted (Charles and Finan 1991), complementation would require a lactose transporter be encoded within the overlapping region (Fig. S1B). We detected an ABC-type transporter system consisting of periplasmic solute-binding protein, permease, and ATP-binding protein (ORF19-ORF17; GenBank AGW45491-AGW45493), but the transporter is probably involved in metal ion uptake. However, ORF21 (GenBank AGW45496) is predicted to encode a major facilitator transporter (IPR020846) with 14 transmembrane helices. This protein belonging to the same major facilitator superfamily as *E. coli* lactose permease LacY might be functional as a lactose transporter when expressed in *E. coli* DH5α and *S. meliloti* RmF728.

Lac^+^ clones Lac20, Lac71, and Lac172 isolated in *S. meliloti* shared a region of 14,707 bp (Fig. S3A and S3B), and was 93% identical to a segment (positions 2,578,724 - 2,593,427) of the *S. marcescens* WW4 chromosome (GenBank accession CP003959). The 14 annotated ORFs within this region exhibited 85-100% amino acid sequence identities to the clustered orthologs (Fig. S3B). These data suggest that the cloned DNA in Lac20, Lac71, and Lac172 originated from *γ-Proteobacteria*. One of the two major facilitator transporters (Lac20_ORF31, GenBank accession AHN97675; Lac20_ORF33, GenBank accession AHN97677) might be involved in lactose uptake in *S. meliloti*. We were unable to identify an ORF encoding a known β-galactosidase based on protein sequences.

Examination of the annotated ORFs of the other 13 Lac^+^ cosmids from *S. meliloti* (Table 2) also did not suggest any candidate that resembled known β-galactosidases. Based on a protein sequence comparison to the CAZy database, which showed low level similarity to proteins carrying known GHs (but not β-galactosidases), we chose Lac36W_ORF11 (GenBank accession AGW45517), Lac161_ORF7 (GenBank accession AGW45552), and Lac161_ORF10 (GenBank accession AGW45555) for further analysis of putative β-galactosidase activity. We amplified the selected ORFs with PCR, and cloned the amplicons into expression vectors, generating C-terminal His tags for overexpression in *E. coli* and subsequent affinity purification. Following processing, the resulting affinity-purified proteins were assayed for β-galactosidase activity. Here, we report the biochemical properties of gene products from these three ORFs and confirm their activities on lactose as substrate.

### Biochemical characterization of Lac36W_ORF11

Cosmid Lac36W exhibited β-galactosidase activity in *S. meliloti* (Table S2), but not in *E. coli.* The cosmid contained a metagenomic DNA fragment of 34,259 bp with 67.3% GC (GenBank accession KF255993). The cloned DNA was assigned taxonomically to *Xanthomonas* of the *γ-Proteobacteria* (Table 2).

Protein sequence searches of the predicted 33 ORFs against the CAZy database suggested that Lac36W_ORF11 (GenBank accession AGW45517) showed sequence similarity to the protein ERE_21070 of *Eubacterium rectale* M104/1 (GenBank accession CBK94002), which has three domains: PBP1_LacI_sugar_binding_like, GGDEF (PF00990), and Glyco_hydro_53 (endo-β-1,4-galactanase, PF07745). The Lac36W_ORF1 1 protein has an N-terminal signal peptide of 21 amino acids predicted by signalP 4.0 (Petersen et al. 2011) and two domains (Fig. 1): 7TMR-DISM-7TM (PF07695) of bacterial membrane-associated receptors and diguanylate cyclase (DGC) or GGDEF domain (PF00990). We predict that the 7TMR-DISM-7TM domain might function as a lactose binding domain like other 7TM-containing proteins (Anantharaman and Aravind 2003) and the C-terminal region may act as a β-galactosidase, though it exhibited no similarity to the endo-β-1,4-galactanase domain in the protein ERE_21070 and other known GH family members. Therefore, we cloned the entire Lac36W_ORF11 and expressed it in *E. coli.* Purified Lac36W_ORF11 protein was able to hydrolyze lactose to galactose and glucose (Table 3). The enzyme maintained 75% activity in the pH range of 6.5 - 8.0 (Fig. 2A) and still kept 20% activity at 50 C (Fig. 2B). Because there was no similarity to any known GH domain and carbohydrate binding module (CBM), we proposed that Lac36W_ORF11 (GenBank, AGW45517) is a new β-galactosidase with possible other functions.

**Figure 2.**
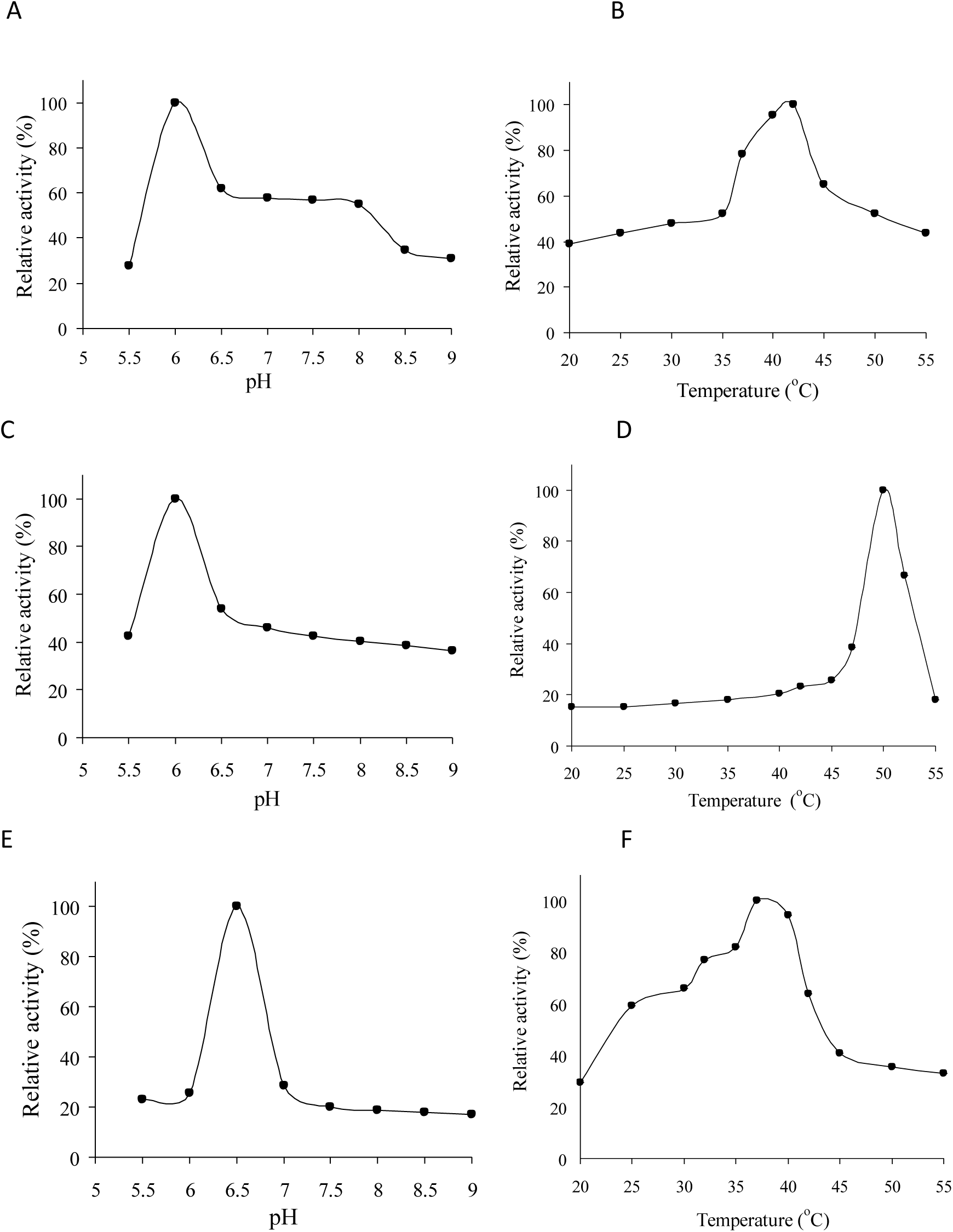
Biochemical characterization of novel β-galactosidases. pH profiles of Lac36W_ORF11 (A), Lac161_ORF7 (C), Lac161_ORF10 (E). Temperature profiles of Lac36W_ORF11 (B), Lac161_ORF7 (D), Lac161_ORF10 (F).

**Table 3.** Biochemical characterization of novel β-galactosidases from 12AC Lac^+^ metagenomic clones complementing *S. meliloti* RmF728 (*lac*)

| ORFs | Proteins | Molecular weight (kDa) | pI | $K_m$ (mM) | $K_{cat}$ (s<sup>-1</sup>) | $K_{cat}/K_m$ (s<sup>-1</sup> M<sup>-1</sup>) | Temperature (°C) | pH |
| --- | --- | --- | --- | --- | --- | --- | --- | --- |
| Lac36W_ORF11 | β-Galactosidase (GenBank, AGW45517) | 79.6 | 8.2 | 2.5 | 10.4 | $3.7 \times 10^3$ | 42 | 6.0 |
| Lac161_ORF7 | β-Galactosidase (GenBank, AGW45552) | 109.0 | 6.7 | 1.8 | 13.2 | $7.3 \times 10^4$ | 50 | 6.0 |
| Lac161_ORF10 | β-Galactosidase (GenBank, AGW45555) | 63.7 | 9.1 | 3.2 | 8.6 | $2.7 \times 10^3$ | 37 | 6.5 |

The Lac36W_ORF1 1 was situated within a putative operon, flanked by Lac36W_ORF12, immediately downstream, and Lac36W_ORF10, immediately upstream. Lac36W_ORF12 encodes a putative methionine-S-sulfoxide reductase and is located 107-bp downstream of the Lac36W_ORF11 (Fig. S4A), whereas Lac36W_ORF10, encoding a hypothetical protein (DUF2007), was located 5 bp upstream of Lac36W_ORF11. The nature of the promoter for this predicted operon and its basis for function in *S. meliloti*, but not *E. coli*, is not known. The Lac36W_ORF14 is predicted to encode a transcriptional regulator (LysR-like) but whether it has a role in regulation of the operon is unknown. Additionally, there were no ORFs encoding homologs to known transporters in the cloned 34-kb DNA. Thus, uptake of lactose and regulation of the predicted operon are unknown.

### Biochemical characterization of Lac161_ORF7

Cosmid Lac161 complemented the Lac^-^ phenotype of *S. meliloti* RmF728 but could not complement *E. coli* DH5α. The cosmid carried an insert of 35,906 bp with 59.1% GC content (GenBank accession KF255994). The metagenomic DNA was assigned taxonomically to *Chthoniobacter* of the phylum *Verrucomicrobia* (Table 2).

Among the annotated 29 ORFs, Lac161_ORF7 (GenBank accession AGW45552) was predicted to be a membrane-bound dehydrogenase protein with three domains (Fig. 1): Piru-Ver-Nterm (TIGR02604), Heat repeat 2 (PF13646), and a cytochrom_C (PF00034) with a putative heme-binding motif CxxCH (TIGER02603). The Heat_2 and Cytochrom_C domains might be involved in intracellular transport and electron transfer. In addition, Lac161_ORF7 was homologous to several proteins annotated as probable glycoside hydrolases, such as HVO_B0215 (GenBank accession ADE01485.1; CBM16, CAZy) of *Haloferax volcanii* DS2. Further sequence alignment analysis did not show any similarity to known GH and CBM. To determine whether the gene product exhibited any GH activity, the Lac161_ORF7 was cloned and expressed. Purified ORF7 protein was able to hydrolyze lactose with a *K*_*m*_ of 1.8 mM, which is the lowest of the three β-galactosidases studied in this work (Table 3). In addition, the *K*_*m*_ value of Lac161_ORF7 is similar to the reported *K*_*m*_ (2.0) of *E. coli* LacZ (Wallenfels and Malhotra 1961). The ORF7 protein was most active at the same pH of 6.0 as Lac36W_ORF11 (Table 3; Fig. 2A and 2C), but the highest activity of Lac161_ORF7 was observed at 50°C (Fig. 2D). In addition, Lac161_ORF7 had the highest *K*_*cat*_/*K*_*m*_ among the β-galactosidases identified in this study. These results implied that Lac161_ORF7 (GenBank accession AGW45552) is the first reported member of a novel β-galactosidase family.

### Biochemical characterization of Lac161_ORF10

Protein sequence comparison with the Pfam database suggested that Lac161_ORF10 (GenBank accession AGW45555) grouped to a family of proteins of unknown function DUF377 (PF04041; Fig. 1), some of which are predicted to be β-fructosidases (GH32 and GH68), and α-L-arabinase and β-xylosidase (GH43 and GH62) (Naumoff 2001). Because of this observation, the Lac161_ORF10 was overexpressed and purified. The resulting gene product was able to hydrolyze lactose with a *K*_*m*_ of 3.2, similar to the values of Lac36W_ORF11 and Lac161_ORF7 (Table 3). The optimal pH and temperature of β-galactosidase activity was 6.5 and 37°C, respectively (Fig. 2E and 3F). In order to further investigate the range of substrate specificity, four other disaccharides were tested as substrates. When sucrose (glucose-β-1,2-fructose) was added, no glucose was released, suggesting that Lac161_ORF10 was not a β-fructofranosidase (or invertase, GH32). Additionally, the ORF10 protein was unable to catalyze hydrolysis of xyloside (xylose-β-1,4-xylose, often associated with GH43), maltose (glucose-β-1,4-glucoside, often associated with GH65), and cellobiose (glucose-α-1,4-glucoside, often associated with GH1). Sequence analysis and activity assays therefore suggested that Lac161_ORF10 (GenBank accession AGW45555) is also a new β-galactosidase, like Lac36W_ORF11 (GenBank accession AGW45517) and Lac161_ORF7 (GenBank accession AGW45552) proteins.

Lac161_ORF7 and ORF_10 encoding the two novel β-galactosidases might form one operon along with Lac161_ORF8 and Lac161_ORF9 (Fig. S4B). The Lac161_ORF8 encodes a hypothetical protein (GenBank accession AGW45553) homologous to an enolase superfamily including o-succinylbenzoate synthase (cd03320). Lac161_ORF9 encodes a hypothetical protein (GenBank accession AGW45554) with a similar domain to methane oxygenase (PF14100). The reason for gene expression in *S. meliloti* but not *E. coli* is not yet known.

### Bioinformatic and structural modeling of Lac161_ORF10

A search of Lac161_ORF10 (GenBank accession AGW45555) against the NCBI Conserved Domain Database (CDD, (Marchler-Bauer et al. 2013)) revealed no significant hits to characterized protein domains (*E* < 0.01). However, a glycoside hydrolase superfamily domain (GH43_62_32_68 superfamily, cl14647) was detected over region 130-234 as the top-scoring CDD hit overall (*E* = 0.10). More specifically, the match corresponds to a GH_J clan domain, which includes GH32 and GH68 enzymes. The presence of a GH_J domain within Lac161_ORF10 is further supported by the domain architectures of related sequences. The top 10 homologs of Lac161_ORF10 detected by BLAST were mainly from *Bacteroides* (Table S3), and all possessed this domain over the aligning region (*E* < 0.01). According to CDTree, Lac161_ORF10 represented a highly distinct branch within the GH_J sequence cluster (Fig. 3A), which provided some explanation for the observed weak similarity to existing CDD domains.

**Figure 3.**
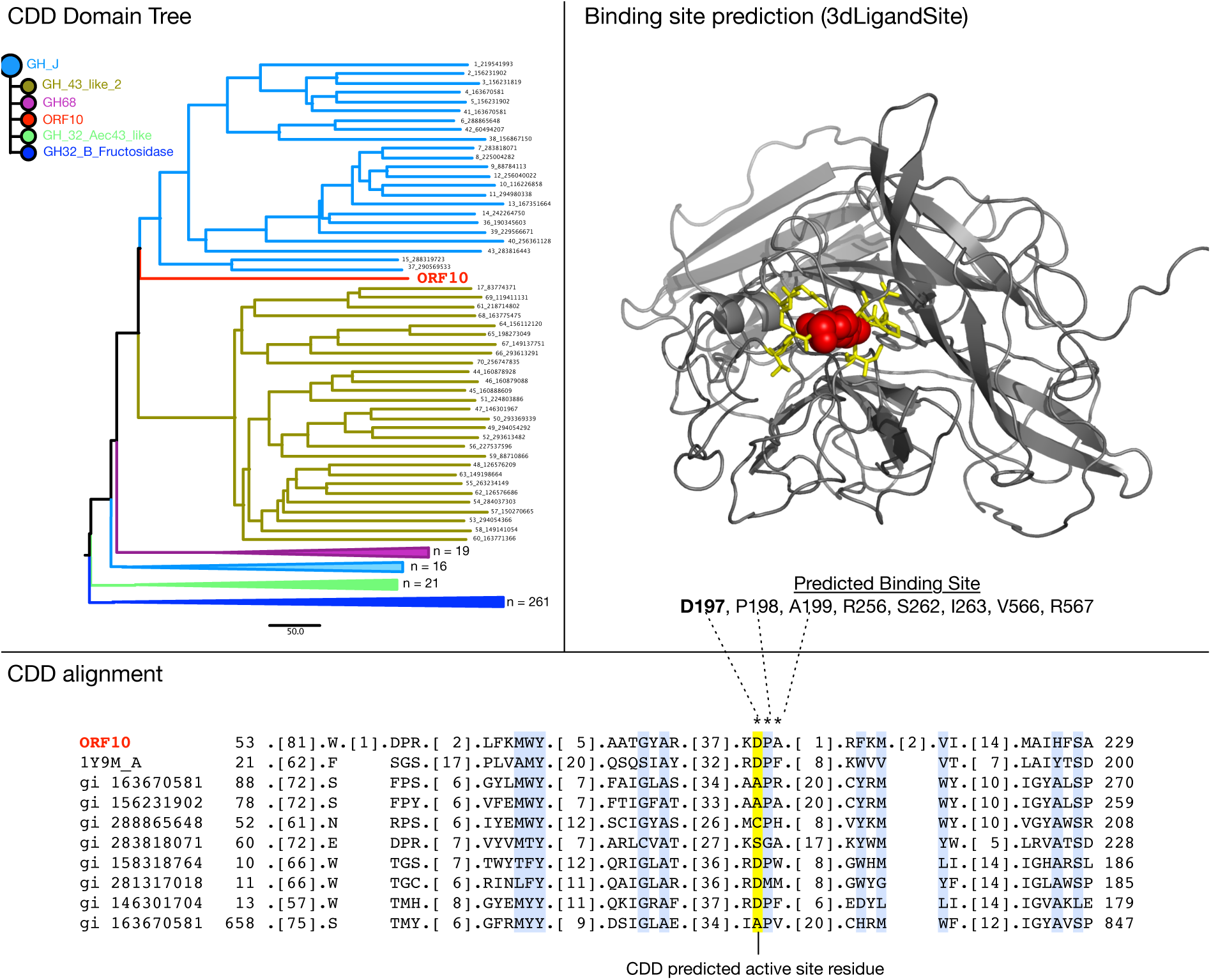
Bioinformatic characterization of a putative glycosyl hydrolase domain in Lac161_ORF10. (A) The NCBI Conserved Domain Database (CDD) predicts Lac161_ORF10 as a divergent member of the GH_J clan of glycosyl hydrolases. (B) Structural model of Lac161_ORF10 generated by Phyre 2.0, with a predicted cluster of 8 Ligand-binding residues highLighted in yellow. The putative binding site was predicted by 3dLigandSite based on the Phyre model with PDB ID 1vkd (chain A) as the template. A NAG Ligand is shown in red, which approximates the location of a lactose molecule in Lac161_ORF10. (C) An alignment of Lac161_ORF10 with the most similar members of the CDD’s GH_J sequence cluster (NCBI gi accession #s are included on the right of the tree). The most conserved columns are coloured Light blue. A predicted active site feature (D197) is highLighted in yellow, and is consistent with 3dLigandSite’s predicted cluster of Ligand-binding residues.

Proteins within the GH_J superfamily, including GH32 and GH68, all posses a five-bladed propeller fold, and share a funnel-shaped active site typically composed of a catalytic nucleophile (e.g., Asp) and proton donor (e.g., Glu) acting as the general acid/base as well as a RDP motif (Lammens et al. 2009) involved in stabilizing the transition state (Fig. 3B). Our analysis suggests that Lac161_ORF10 also shared some of these characteristics.

Using Phyre 2.0 (Kelley and Sternberg 2009), a structural model of Lac161_ORF10 was generated. Phyre predicted a five-bladed propeller fold for Lac161_ORF10 (Fig. 3B) with high confidence (99.9%) based on the template PDB 1vkd_A, a predicted glycoside hydrolase from *Thermotoga maritima* (Tmari_1232). Interestingly, both Tmari_1232 and Lac161_ORF10 are members of the Pfam DUF377 family, further supporting the model. We then analyzed potential active sites using two separate methods: a sequence and structure-based approach. According to the CDD sequence alignment, Lac161_ORF10 possesses a KDP motif (residues 196-198) that aligns to the active site RDP motif in the reference 1y9m structure (Fig. 3C). Ligand-binding sites were also predicted in the structural model using 3dLigandSite (Wass et al. 2010). This revealed a predicted cluster of eight residues, including the previously identified D-197 residue, as forming the putative active site (Fig. 3B). However, alternate alignments and putative active sites from those reported above are possible given the structural repetition of five-bladed propellers. Ultimately, Lac161_ORF10 (GenBank, AGW45555) appears to represent a novel family of β-galactosidase with a GH_J-like five-bladed propeller glycoside hydrolase domain, and an active site similar in composition to other members of this superfamily.

### Metagenome abundance

We were interested in the distribution of sequences similar to the newly described β-galactosidase sequences throughout different metagenomes. To address this, we performed protein homology searches with these sequences against collections of aquatic, human gut and soil metagenomic databases, and normalized using the housekeeping *rpoB* abundance (Figure 4). Homologs to each of the three genes are represented in all three habitats. However, Lac36W_ORF11 in human gut is by far of greatest relative abundance. Lac36W_ORF11 is also high in soil, but not as high as in human gut. Although of overall lower relative abundance, Lac161_ORF10 is also of greater abundance in human gut than in soil or aquatic. Lac161_ORF7 exhibits a quite different profile, being extremely rare in human gut, low levels in aquatic, but higher levels in soil. It will be of interest to determine whether these homologs are also functional β-galactosidases.

**Figure 4.**
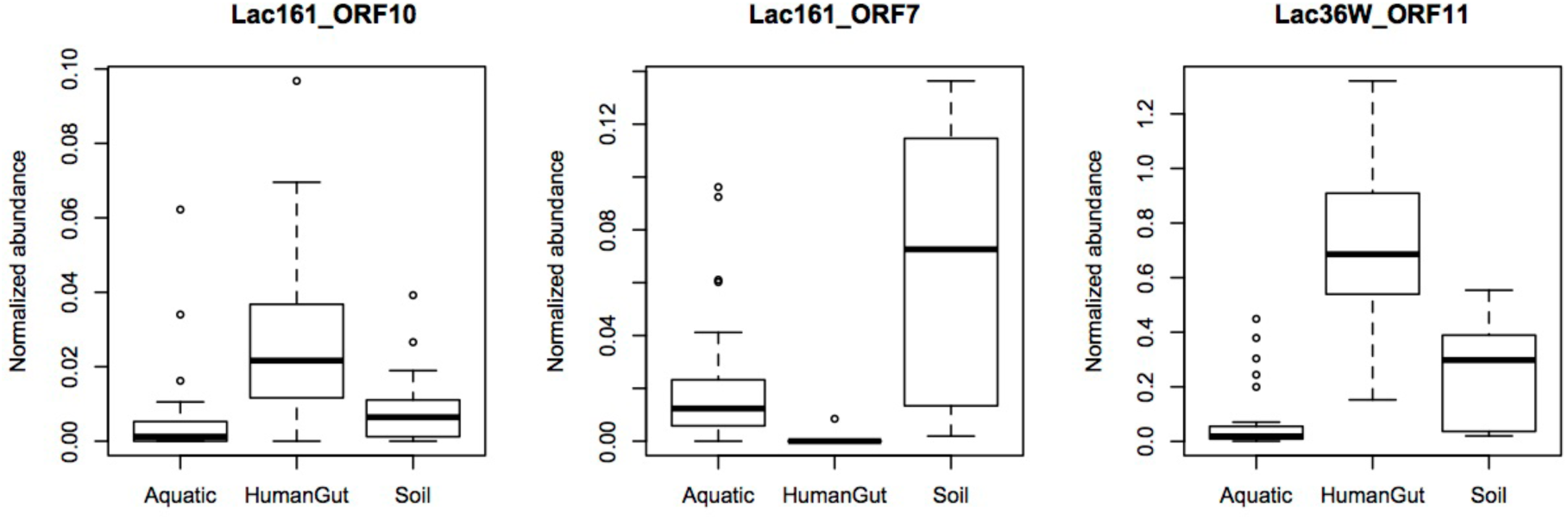
Protein homology searches of novel β-galactosidase sequences of Lac161_ORF10, Lac161_ORF7 and Lac36W_ORF1 1 against aquatic, human gut and soil metagenomic databases, normalized to the housekeeping *rpoB* gene.

### Host influence on screening

By discovering founding members of three novel *β*-galactosidase families, we have reinforced the value of functional metagenomics for isolating novel genes that could not have been predicted from DNA sequence analysis alone. Activity-based screening of metagenomic library clones for biocatalysts is dependent on the expression of genes of interest and presence of accessory components required for the enzyme activity in the surrogate hosts (Martinez et al. 2004; Taupp et al. 2011). Multi-host-systems have been developed to improve functional screening (Aakvik et al. 2009; Biver et al. 2013; Craig et al. 2010; Li et al. 2005; Ly et al. 2011; Wang et al. 2006; Wexler and Johnston 2010). In the present work, functional screening of the corn field soil library (12AC) for the ability to complement β-galactosidase mutants resulted in a greater number of distinct clones using *S. meliloti* than the most widely used *E. coli.* In addition, three novel β-galactosidase genes were identified only in *S. meliloti*. These data emphasize the indispensable development of multi-host systems for functional screening.

## Discussion

Metagenomics provides unprecedented access to the genomic potential of uncultivated microbial communities. Despite enormous progress resulting from developments in high throughput sequencing, the potential for novel enzyme discovery remains highest using a functional metagenomics approach, in which genes are isolated based on their function rather than by DNA sequence similarity to already known genes. Using such an approach, we have discovered genes encoding novel types of lactose hydrolyzing enzymes. The enzymes encoded by these genes were biochemically similar to known enzymes, although they would not have been easily predicted by their sequences without knowing that they were carried on a segment of DNA that encoded β-galactosidase activity. These results demonstrate the importance of sequence-agnostic functional screens for the discovery of enzymes of novel origin, and suggest that further implementation of this strategy will contribute to fundamental knowledge about the relationship between sequence and protein function, improve the resolution of sequence based metagenomics, and expand the repertoire of novel enzymes available for industrial applications.

This work follows on other metagenomic functional screening efforts that have discovered β-galactosidases of GH1 (Gupta et al. 2012), GH42 (Wang et al. 2010) (Zhang et al. 2013), GH43 (Ferrer et al. 2012; Wierzbicka-Wos et al. 2013), and two new GH members (Beloqui et al. 2010). Here we have highlighted the application of functional metagenomics for mining novel enzymes from soil microbial communities/ While the functional metagenomics strategy has potential for expanding the availability of enzymes that can be further developed for biotech applications, it is perhaps just as important to apply such strategies to the expansion of knowledge that will inform functional interpretation of DNA sequence. This in turn could impact on the ability to derive metabolic information from genome sequence, even from uncultivated organism. We suggest that the use of a diversity of surrogate hosts for functional metagenomic screening has the potential to substantially extend the breadth of gene discovery.

## Nucleotide sequence accession numbers

Complete sequences of metagenomic Lac^+^ cosmids have been deposited in GenBank (Table 2), accession numbers: KF255992-KF255994, KF796593-KF796611

## Abbreviations

Cm^R^: (chloramphenicol resistant)
Gm^R^: (gentamicin resistant)
Km^R^: (kanamycin resistant)
Nm^R^: (neomycin resistant)
Rif^R^: (rifampicin resistant)
Sm^R^: (streptomycin resistant)
Tc^R^: (tetracycline resistant)

## Acknowledgements

We are grateful to Julia Hanchard and Shirley Wong for technical assistance.

## Funding

This work was financially supported by a Strategic Projects grant and Discovery Grants from the Natural Sciences and Engineering Research Council of Canada (NSERC).

## Conflict of interest

The authors declare no conflict of interest.

## Ethical statement

The authors certify that this manuscript has not been published previously, and not under consideration for publication elsewhere, in whole or in part. No data have been fabricated or manipulated (including images), and no data, text, or theories by others are presented as if they were the authors’ own. Consent to submit has been received explicitly from all the authors listed, and authors whose names appear on the submission have contributed sufficiently to the scientific work and therefore share collective responsibility and accountability for the results. This article does not contain any studies with human participants or animals performed by any of the authors.

## Supplementary tables

**Table S1.**
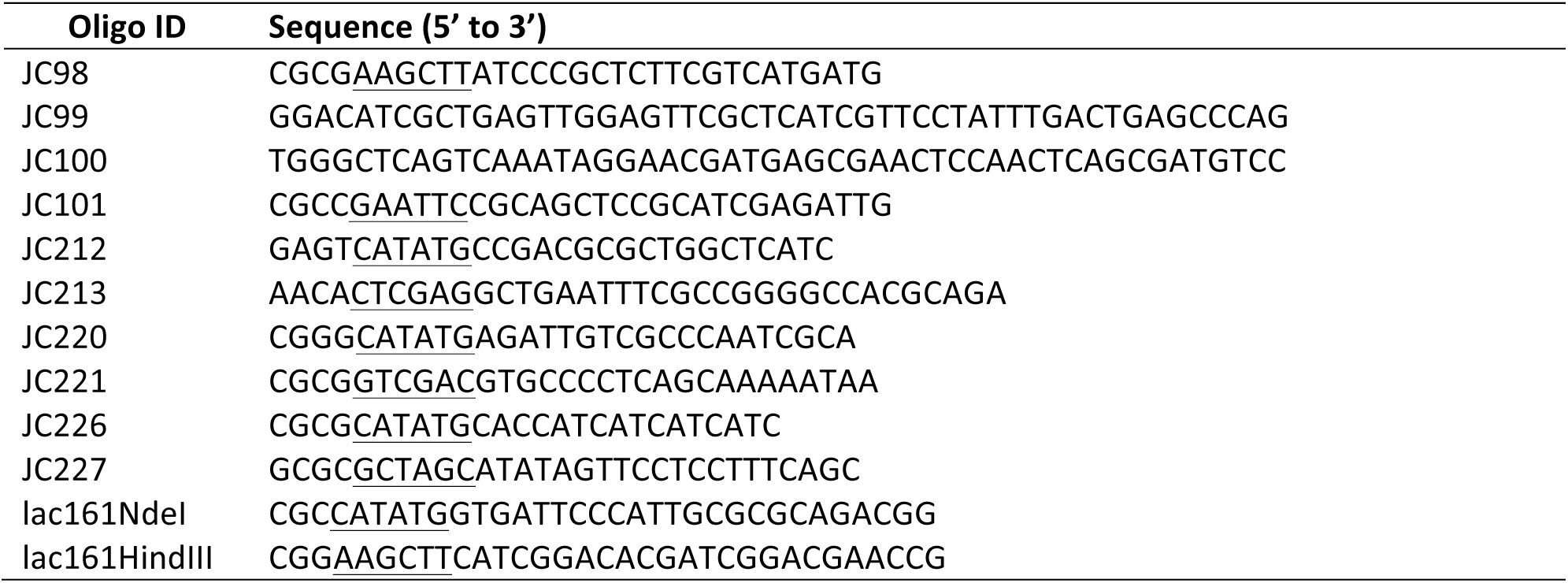
DNA oLigonucleotides used in this study with restriction recognition sites underlined.

**Table S2.** β-Galactosidase activities of random 12AC *lac^+^* clones in *S. meliloti* SmUW253 *(lacZ1).* Cosmid pJC8 was used as a negative control. Clone numbers in bold font also complemented *E. coli* DH5α (Rif^R^) in M9-lactose medium. Underlined clone numbers were completely sequenced.

| 12AC *lac*<sup>+</sup> clone | β-Galactosidase specific activity ± standard deviation |
| --- | --- |
| **12** | 9902 ± 1052 |
| <u>16</u> | 4379 ± 643 |
| **34** | 15596 ± 1633 |
| 55 | 4388 ± 260 |
| 77 | 3958 ± 700 |
| 78 | 4301 ± 814 |
| 86 | 3226 ± 470 |
| 94 | 3292 ± 610 |
| 99 | 3671 ± 626 |
| 114 | 3558 ± 123 |
| 115 | 3240 ± 212 |
| 128 | 3596 ± 508 |
| <u>160</u> | 3094 ± 15 |
| <u>161</u> | 3916 ± 174 |
| 169 | 3606 ± 433 |
| <u>172</u> | 3903 ± 327 |
| **177** | 15445 ± 796 |
| 183 | 3331 ± 440 |
| 199 | 3851 ± 155 |
| 204 | 3357 ± 140 |
| 234 | 3589 ± 352 |
| 253 | 1801 ± 177 |
| 266 | 4001 ± 343 |
| 270 | 4351 ± 717 |
| 271 | 3835 ± 344 |
| 305 | 3679 ± 52 |
| 306 | 3770 ± 107 |
| 319 | 3788 ± 408 |
| 150B | 3544 ± 633 |
| 150W | 3889 ± 545 |
| 175B | 4163 ± 582 |
| 195W | 4211 ± 1099 |
| 203W | 4112 ± 251 |
| **206B** | 15598 ± 1104 |
| 206W | 4147 ± 309 |
| <u>24B</u> | 16973 ± 1348 |
| 267A | 3707 ± 468 |
| **35B** | 16647 ± 664 |
| <u>36W</u> | 3772 ± 310 |

**Table S3.** Top twenty homologs of Lac161_ORF10 detected by a BlastP search of the NCBI nr database.

| Species | Protein (GenBank accession #) | Function | % identity | E-value | Bit score |
| --- | --- | --- | --- | --- | --- |
| *Bacteroides* sp. D22 | gi:496330437 | Conserved hypothetical protein | 44.9 | 1.00E-145 | 444 |
| *Capnocytophaga* sp. oral taxon 329 str. F0087 | gi:725815884 | Hypothetical protein | 45.8 | 1.00E-145 | 444 |
| *Bacteroides finegoldii* | gi:515710157 | Hypothetical protein | 44.5 | 2.00E-145 | 443 |
| *Bacteroides finegoldii* CL09T03C10 | gi:408473640 | Hypothetical protein | 44.5 | 4.00E-145 | 442 |
| *Chthoniobacter flavus* | gi:494038897 | Glycosyl hydrolase family 32 domain protein | 47.3 | 8.00E-145 | 442 |
| *Bacteroides* (multiple species) | gi:490423864 | Hypothetical protein | 44.1 | 1.00E-143 | 439 |
| *Bacteroides* sp. 2_2_4 | gi:229449608 | Hypothetical protein | 44.4 | 2.00E-143 | 438 |
| *Capnocytophaga* sp. oral taxon 324 | gi:496921710 | Hypothetical protein | 45.1 | 4.00E-143 | 437 |
| *Capnocytophaga* sp. oral taxon 329 str. F0087 | gi:725815883 | Hypothetical protein | 44.9 | 6.00E-143 | 437 |
| *Capnocytophaga* sp. oral taxon 324 | gi:496921709 | Hypothetical protein | 44.4 | 2.00E-141 | 434 |
| *Bacteroides eggerthii* | gi:490420689 | Hypothetical protein | 43.1 | 5.00E-139 | 427 |
| *Capnocytophaga* sp. oral taxon 326 | gi:496931704 | Hypothetical protein | 43.9 | 2.00E-138 | 425 |
| *Bacteroides eggerthii* CAG:109 | gi:547198227 | Conserved hypothetical protein | 43.0 | 2.00E-138 | 425 |
| *Bacteroides* (multiple species) | gi:496046067 | Tat pathway signal sequence domain protein | 44.4 | 3.00E-138 | 426 |
| *Bacteroides eggerthii* | gi:490420690 | Uncharacterized protein | 44.7 | 1.00E-136 | 421 |
| *Bacteroides eggerthii* CAG:109 | gi:547198228 | Uncharacterized protein | 44.7 | 2.00E-136 | 421 |
| *Bacteroides* sp. 3_1_23 | gi:495921857 | Conserved hypothetical protein | 41.9 | 8.00E-135 | 417 |
| *Bacteroides eggerthii* | gi:490418291 | Hypothetical protein | 43.0 | 5.00E-133 | 411 |
| *Bacteroides* sp. 2_1_22 | gi:262356909 | Hypothetical protein | 44.0 | 5.00E-132 | 410 |
| *Bacteroides xylanisolvens* SD CC 2a | gi:292638199 | Conserved hypothetical protein | 44.0 | 7.00E-132 | 409 |

**Table S4.** Detected abundance of three novel beta-galactosidases in a variety of metagenomic datasets.

| BIOME | DATASET | Lac161_10 | Lac161_ORF7 | Lac36W_ORF11 | RPOB |
| --- | --- | --- | --- | --- | --- |
| Aquatic | CAM_P_0000692_MILOCO.orfs.fa | 92 | 3443 | 3033 | 191475 |
| Aquatic | Antarctica_Aquatic_Microbial_Metagenome.orfs.fa | 208 | 1629 | 10066 | 151451 |
| Aquatic | CAM_PROJ_GOS.orfs.fa | 65 | 921 | 3262 | 61673 |
| Aquatic | CAM_PROJ_YLake.orfs.fa | 69 | 987 | 2678 | 47527 |
| Aquatic | CAM_P_0001026_LagrangianSamplingMontereyBay.orfs.fa | 7 | 127 | 34 | 33957 |
| Aquatic | ALOHA_orfs.fa | 54 | 175 | 170 | 12165 |
| Aquatic | CAM_PROJ_WesternChannelOMM.orfs.fa | 10 | 129 | 452 | 10353 |
| Aquatic | CAM_PROJ_BATS.orfs.fa | 91 | 268 | 99 | 10349 |
| Aquatic | CAM_P_0000719_MontereyBayTransect.orfs.fa | 15 | 76 | 35 | 9893 |
| Aquatic | CAM_PROJ_HOT_pyrosequenced.orfs.fa | 17 | 93 | 104 | 8525 |
| Aquatic | CAM_P_0001136_MahoneyLake.orfs.fa | 209 | 179 | 2328 | 6145 |
| Aquatic | CAM_P_0000545_Guaymas_Basin.orfs.fa | 14 | 307 | 360 | 5098 |
| Aquatic | CAM_P_0001028_NorthPacificMetagenomes.orfs.fa | 14 | 1 | 24 | 4920 |
| Aquatic | CAM_P_0001129_DeepChlorophyllMaximum.orfs.fa | 2 | 35 | 13 | 4283 |
| Aquatic | CAM_P_0000712_Bermuda_Oceanic.orfs.fa | 69 | 175 | 70 | 4254 |
| Aquatic | CAM_P_0001130_SantaPolaSaltern.orfs.fa | 0 | 0 | 142 | 3995 |
| Aquatic | CAM_PROJ_HOT.orfs.fa | 39 | 95 | 77 | 3701 |
| Aquatic | CAM_PROJ_Bacterioplankton.transeq.ORFs.fa | 2 | 44 | 17 | 3586 |
| Aquatic | CAM_PROJ_MontereyBay.orfs.fa | 1 | 27 | 41 | 3286 |
| Aquatic | CAM_PROJ_PacificOcean.orfs.fa | 0 | 16 | 16 | 2628 |
| Aquatic | CAM_PROJ_PML.orfs.fa | 0 | 3 | 69 | 2480 |
| Aquatic | CAM_PROJ_Sapelo2008.orfs.fa | 0 | 28 | 28 | 2206 |
| Aquatic | CAM_PROJ_AmazonRiverPlume.orfs.fa | 0 | 6 | 22 | 2176 |
| Aquatic | CAM_PROJ_GeneExpression.orfs.fa | 0 | 6 | 17 | 1973 |
| Aquatic | CAM_PROJ_BisonMetagenome.orfs.fa | 3 | 9 | 496 | 1633 |
| Aquatic | CAM_PROJ_IceMetagenome.orfs.fa | 0 | 29 | 367 | 1503 |
| Aquatic | CAM_P_0001133_HypersalineCoastalLagoon.orfs.fa | 0 | 9 | 72 | 1415 |
| Aquatic | Polar_Metagenome.orfs.fa | 8 | 121 | 71 | 1309 |
| Aquatic | CAM_P_0001132_FreshwaterLagoon.orfs.fa | 10 | 67 | 219 | 1097 |
| Aquatic | CAM_P_0001131_SantaPolaSaltern.orfs.fa | 0 | 0 | 1 | 1094 |
| Aquatic | Arctic_seawater_EBI_ERS089005.orfs.fa | 9 | 88 | 46 | 915 |
| Aquatic | CAM_PROJ_HypersalineMat.transeq.ORFs.fa | 14 | 4 | 101 | 225 |
| HumanGut | MH0012.fa | 5 | 0 | 181 | 437 |
| HumanGut | MH0006.fa | 8 | 0 | 280 | 310 |
| HumanGut | MH0009.fa | 5 | 0 | 120 | 255 |
| HumanGut | MH0086.fa | 3 | 0 | 254 | 247 |
| HumanGut | MH0050.fa | 2 | 0 | 91 | 242 |
| HumanGut | MH0082.fa | 8 | 0 | 151 | 239 |
| HumanGut | MH0011.fa | 2 | 0 | 152 | 222 |
| HumanGut | V1.CD-14.fa | 1 | 0 | 137 | 219 |
| HumanGut | MH0040.fa | 7 | 0 | 110 | 217 |
| HumanGut | V1.CD-8.fa | 8 | 0 | 100 | 217 |
| HumanGut | MH0081.fa | 8 | 0 | 125 | 212 |
| HumanGut | MH0069.fa | 3 | 0 | 144 | 211 |
| HumanGut | MH0065.fa | 3 | 0 | 162 | 201 |
| HumanGut | MH0042.fa | 3 | 0 | 94 | 196 |
| HumanGut | MH0070.fa | 5 | 0 | 182 | 196 |
| HumanGut | MH0054.fa | 7 | 0 | 91 | 194 |
| HumanGut | MH0060.fa | 2 | 0 | 78 | 192 |
| HumanGut | V1.CD-4.fa | 2 | 0 | 140 | 190 |
| HumanGut | MH0053.fa | 0 | 0 | 103 | 189 |
| HumanGut | MH0055.fa | 3 | 0 | 161 | 188 |
| HumanGut | MH0003.fa | 5 | 0 | 125 | 187 |
| HumanGut | MH0043.fa | 4 | 0 | 92 | 187 |
| HumanGut | MH0079.fa | 3 | 0 | 29 | 184 |
| HumanGut | V1.UC-19.fa | 1 | 0 | 124 | 183 |
| HumanGut | MH0077.fa | 2 | 0 | 192 | 181 |
| HumanGut | MH0080.fa | 5 | 0 | 167 | 179 |
| HumanGut | MH0059.fa | 3 | 0 | 163 | 178 |
| HumanGut | MH0075.fa | 8 | 0 | 95 | 178 |
| HumanGut | MH0031.fa | 2 | 0 | 109 | 177 |
| HumanGut | MH0066.fa | 7 | 0 | 49 | 175 |
| HumanGut | MH0025.fa | 2 | 0 | 109 | 173 |
| HumanGut | V1.UC-6.fa | 5 | 0 | 183 | 173 |
| HumanGut | MH0063.fa | 2 | 0 | 102 | 171 |
| HumanGut | V1.UC-9.fa | 8 | 0 | 83 | 171 |
| HumanGut | MH0035.fa | 3 | 0 | 150 | 170 |
| HumanGut | MH0083.fa | 6 | 0 | 132 | 170 |
| HumanGut | MH0033.fa | 2 | 0 | 145 | 169 |
| HumanGut | MH0038.fa | 2 | 0 | 76 | 167 |
| HumanGut | MH0014.fa | 8 | 0 | 179 | 165 |
| HumanGut | MH0058.fa | 10 | 0 | 182 | 163 |
| HumanGut | V1.UC-8.fa | 5 | 0 | 124 | 161 |
| HumanGut | MH0030.fa | 2 | 0 | 82 | 160 |
| HumanGut | MH0002.fa | 1 | 0 | 90 | 156 |
| HumanGut | MH0028.fa | 5 | 0 | 127 | 155 |
| HumanGut | MH0064.fa | 15 | 0 | 124 | 155 |
| HumanGut | MH0056.fa | 10 | 0 | 89 | 154 |
| HumanGut | MH0039.fa | 5 | 0 | 146 | 152 |
| HumanGut | V1.CD-13.fa | 8 | 0 | 186 | 151 |
| HumanGut | MH0071.fa | 4 | 0 | 103 | 150 |
| HumanGut | MH0016.fa | 3 | 0 | 155 | 149 |
| HumanGut | MH0041.fa | 3 | 0 | 110 | 148 |
| HumanGut | MH0036.fa | 10 | 0 | 137 | 144 |
| HumanGut | MH0044.fa | 2 | 0 | 167 | 143 |
| HumanGut | V1.CD-9.fa | 1 | 0 | 52 | 142 |
| HumanGut | MH0062.fa | 5 | 0 | 101 | 140 |
| HumanGut | V1.CD-11.fa | 7 | 0 | 91 | 140 |
| HumanGut | MH0020.fa | 2 | 0 | 78 | 139 |
| HumanGut | MH0057.fa | 3 | 0 | 59 | 137 |
| HumanGut | MH0067.fa | 2 | 0 | 129 | 137 |
| HumanGut | MH0052.fa | 0 | 0 | 98 | 136 |
| HumanGut | MH0076.fa | 5 | 0 | 143 | 135 |
| HumanGut | MH0045.fa | 6 | 0 | 173 | 131 |
| HumanGut | MH0068.fa | 3 | 0 | 97 | 126 |
| HumanGut | MH0074.fa | 4 | 0 | 119 | 125 |
| HumanGut | V1.CD-3.fa | 1 | 0 | 86 | 125 |
| HumanGut | MH0026.fa | 1 | 0 | 98 | 123 |
| HumanGut | MH0048.fa | 3 | 0 | 62 | 123 |
| HumanGut | MH0051.fa | 1 | 0 | 78 | 120 |
| HumanGut | MH0061.fa | 7 | 0 | 82 | 120 |
| HumanGut | MH0032.fa | 1 | 1 | 104 | 118 |
| HumanGut | MH0021.fa | 5 | 0 | 108 | 118 |
| HumanGut | MH0037.fa | 5 | 0 | 69 | 118 |
| HumanGut | MH0085.fa | 8 | 0 | 76 | 115 |
| HumanGut | MH0073.fa | 5 | 0 | 142 | 111 |
| HumanGut | MH0046.fa | 4 | 0 | 98 | 109 |
| HumanGut | MH0047.fa | 1 | 0 | 16 | 105 |
| Soil | 4510219.3.transeq.fa | 1001 | 732 | 7714 | 384993 |
| Soil | 4541646.3.transeq.fa | 498 | 833 | 1596 | 74631 |
| Soil | 4541645.3.transeq.fa | 544 | 990 | 1831 | 71208 |
| Soil | 4541647.3.transeq.fa | 474 | 800 | 1783 | 61175 |
| Soil | 4541642.3.transeq.fa | 465 | 771 | 1724 | 60466 |
| Soil | 4541649.3.transeq.fa | 272 | 652 | 1738 | 55281 |
| Soil | 4539063.3.transeq.fa | 343 | 924 | 1515 | 53483 |
| Soil | 4541651.3.transeq.fa | 304 | 557 | 2099 | 51570 |
| Soil | 4541641.3.transeq.fa | 356 | 665 | 1378 | 50646 |
| Soil | 4539064.3.transeq.fa | 393 | 652 | 1545 | 48747 |
| Soil | 4541644.3.transeq.fa | 347 | 623 | 1572 | 47658 |
| Soil | 4541648.3.transeq.fa | 279 | 564 | 1138 | 47348 |
| Soil | 4541650.3.transeq.fa | 305 | 722 | 1765 | 47232 |
| Soil | 4543020.3.transeq.fa | 5 | 52 | 185 | 3671 |
| Soil | 4514245.3.transeq.fa | 4 | 11 | 204 | 3317 |
| Soil | 4543023.3.transeq.fa | 3 | 58 | 172 | 2716 |
| Soil | 4543022.3.transeq.fa | 1 | 24 | 68 | 2490 |
| Soil | 4446153.3.transeq.fa | 22 | 156 | 234 | 1238 |
| Soil | 4543019.3.transeq.fa | 0 | 5 | 66 | 981 |
| Soil | 4543021.3.transeq.fa | 1 | 14 | 36 | 980 |
| Soil | 4537193.3.transeq.fa | 5 | 67 | 317 | 952 |
| Soil | 4537195.3.transeq.fa | 10 | 108 | 262 | 878 |
| Soil | 4537194.3.transeq.fa | 3 | 60 | 177 | 777 |
| Soil | 4537190.3.transeq.fa | 7 | 95 | 249 | 773 |
| Soil | 4537191.3.transeq.fa | 9 | 64 | 146 | 678 |
| Soil | 4537192.3.transeq.fa | 1 | 38 | 177 | 651 |
| Soil | 4478941.3.transeq.fa | 1 | 23 | 95 | 322 |
| Soil | 4478940.3.transeq.fa | 5 | 31 | 131 | 278 |
| Soil | 4478294.3.transeq.fa | 0 | 17 | 87 | 275 |
| Soil | 4478038.3.transeq.fa | 1 | 20 | 81 | 272 |
| Soil | 4479311.3.rhiz.transeq.fa | 3 | 22 | 120 | 271 |
| Soil | 4478937.3.transeq.fa | 2 | 32 | 103 | 255 |
| Soil | 4477790.3.transeq.fa | 2 | 29 | 81 | 253 |
| Soil | 4478290.3.rhiz.transeq.fa | 1 | 29 | 114 | 253 |
| Soil | 4478939.3.rhiz.transeq.fa | 2 | 29 | 105 | 249 |
| Soil | 4477749.3.rhiz.transeq.fa | 0 | 26 | 103 | 243 |
| Soil | 4477751.3.rhiz.transeq.fa | 3 | 23 | 89 | 242 |
| Soil | 4478934.3.rhiz.transeq.fa | 0 | 10 | 93 | 239 |
| Soil | 4478291.3.rhiz.transeq.fa | 3 | 17 | 86 | 237 |
| Soil | 4478222.3.transeq.fa | 0 | 24 | 71 | 234 |
| Soil | 4478283.3.transeq.fa | 0 | 21 | 117 | 232 |
| Soil | 4478030.3.rhiz.transeq.fa | 0 | 20 | 84 | 231 |
| Soil | 4478936.3.transeq.fa | 4 | 26 | 99 | 216 |
| Soil | 4477755.3.rhiz.transeq.fa | 4 | 28 | 99 | 211 |
| Soil | 4478943.3.transeq.fa | 3 | 25 | 97 | 209 |
| Soil | 4477757.3.transeq.fa | 3 | 28 | 73 | 207 |
| Soil | 4478037.3.rhiz.transeq.fa | 0 | 17 | 73 | 206 |
| Soil | 4478938.3.rhiz.transeq.fa | 8 | 15 | 113 | 204 |
| Soil | 4478292.3.rhiz.transeq.fa | 5 | 25 | 103 | 188 |
| Soil | 4477789.3.transeq.fa | 0 | 24 | 53 | 176 |

## Supplementary Figures

**Fig. S1.**
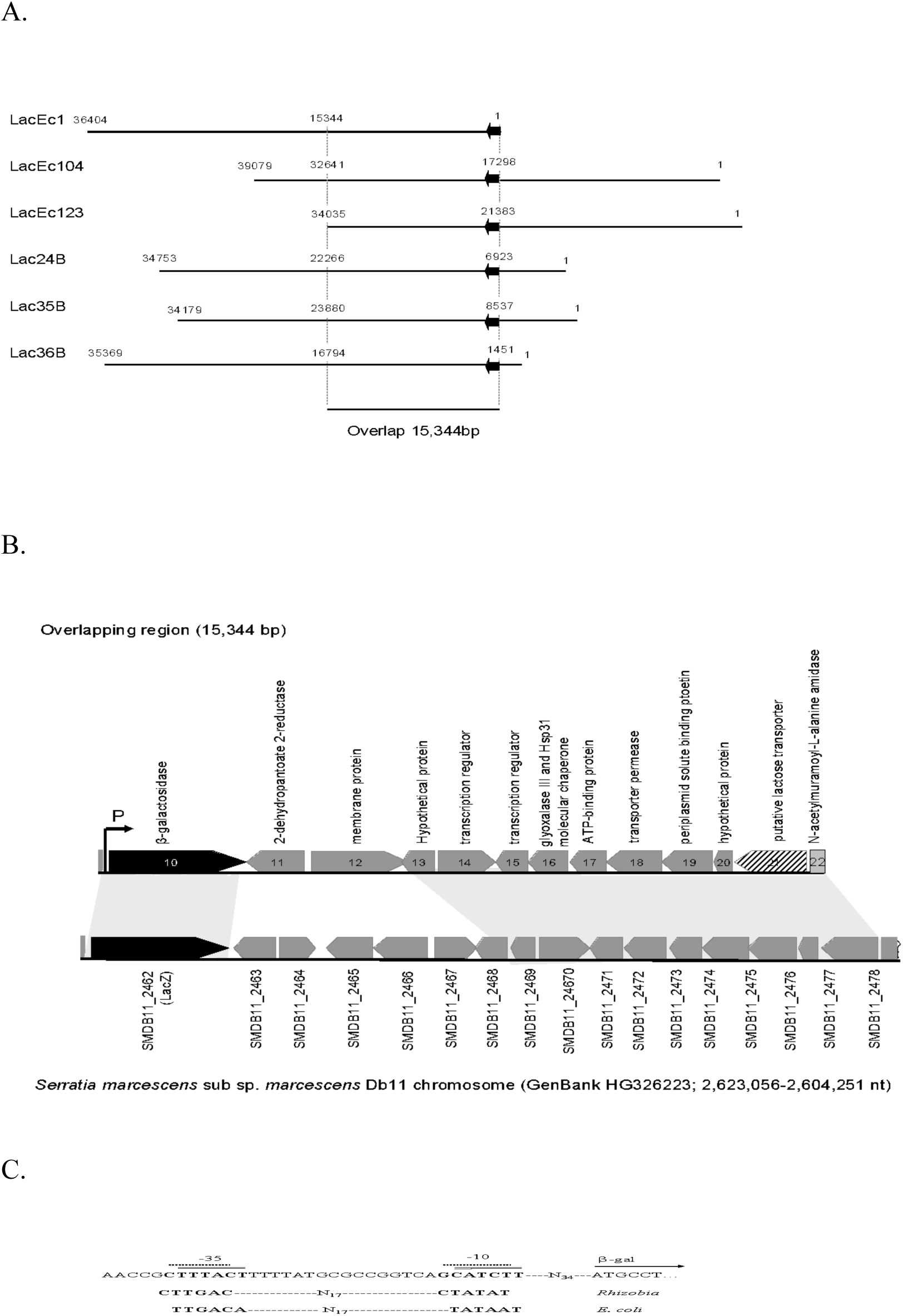
Lac^+^ clones isolated from *E. coli* (LacEc1, LacEc104 and LacEc123) and *S. meliloti* (Lac24B, Lac35B and Lac36B). (A) An overlapping region of 15,344 bp was present in those cosmids. (B) A β-galactosidase of family GH2 (ORF10, solid box), and putative lactose transporter (ORF21, dash lined box) were predicted in Lac35B. The regions encoding orthologs in *γ-Proteobacteria Serratia marcescens* subsp. *marcescens* Db11 chromosome (GenBank HG326223; 2,623,056 - 2,604,251 nt) were highlighted. (C) Putative RpoD promoters (P) active in both *E. coli* and *S. meliloti* were located upstream of the β-galactosidase gene. The same enzyme was encoded by LacEc1_ORF31, LacEc 104ORF20, LacEc123_ORF13, Lac24B_ORF9, and Lac36B_ORF3 respectively.

**Fig. S2.**
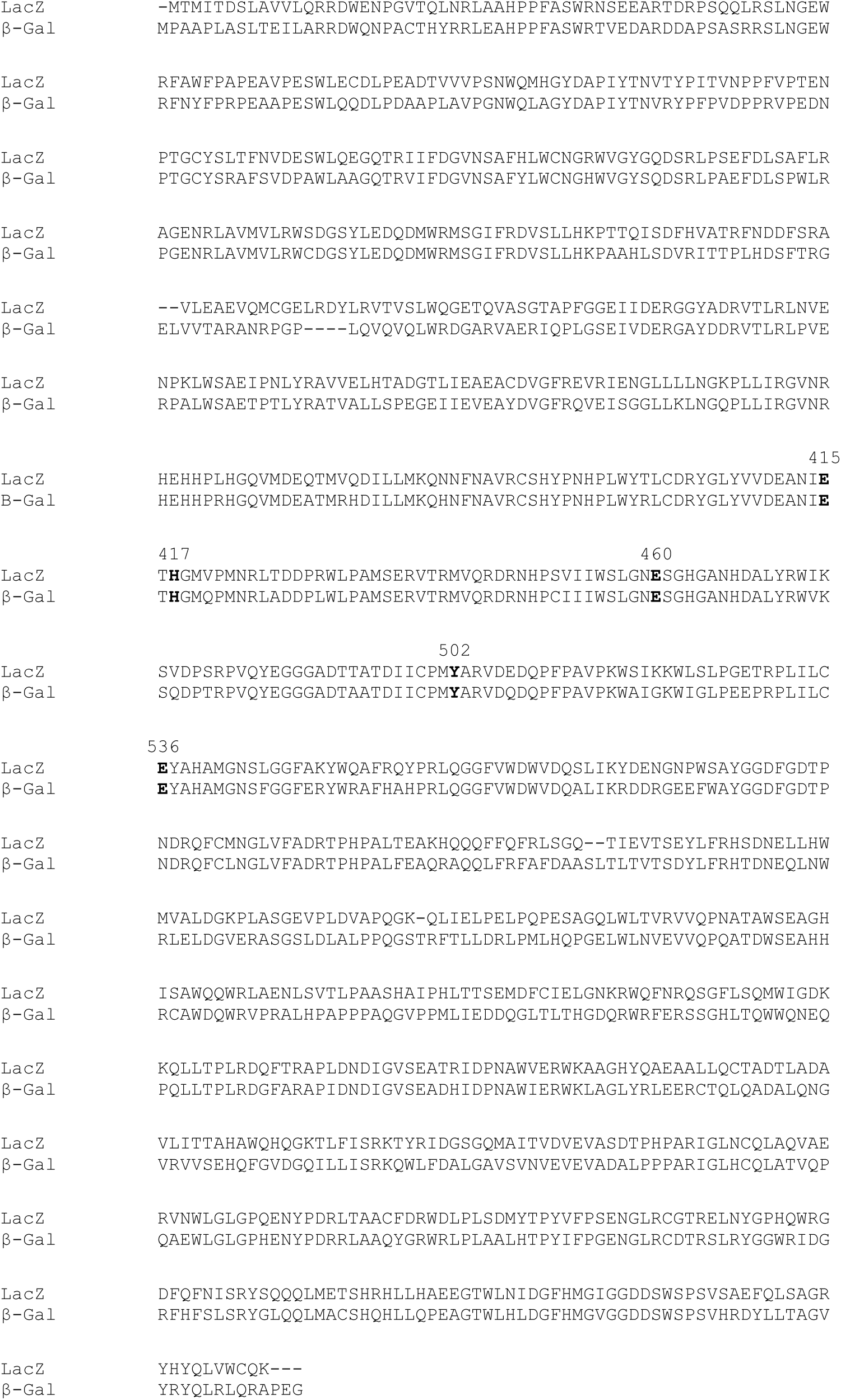
Sequence alignment of β-glactosidases LacZ of *E. coli* K12 substr, W3110 (Genbank, BAE76126) and 12AC metagenomic clone LacEc1 (β-Gal; LacEc1_ORF31; GenBank, KF96609). Conserved amino acids Glu^415^, His^417^, Glu^460^, Tyr^502^ and Glu^536^ at the active sites of LacEc1 ORF31 were highlighted.

**Fig. S3.**
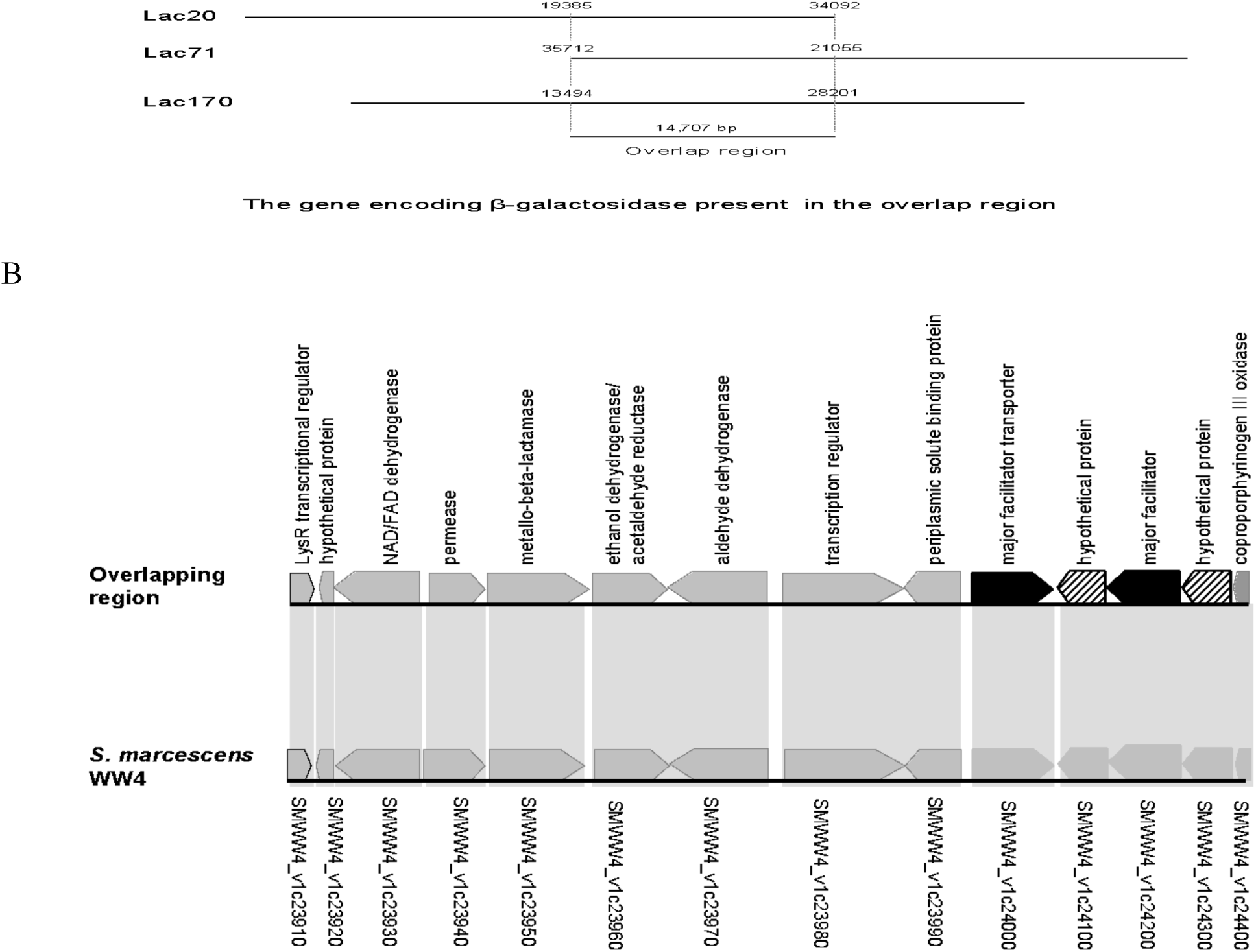
Lac^+^ clone Lac20, Lac71 and Lac172 isolated from *S. meliloti.* (A) An overlapping region of 14,707 bp was present in those cosmids. (B) The major facilitator transporter(s) (solid box) in the region might be involved in lactose uptake. The hypothetical protein(s) (dash lined box) might be a β-galactosidase. Orthologs in *γ-Proteobacteria Serratia marcescens* WW4 chromosome (GenBank CP003959; 2,578,724 - 2,593,247 nt) were highlighted.

**Fig. S4.**
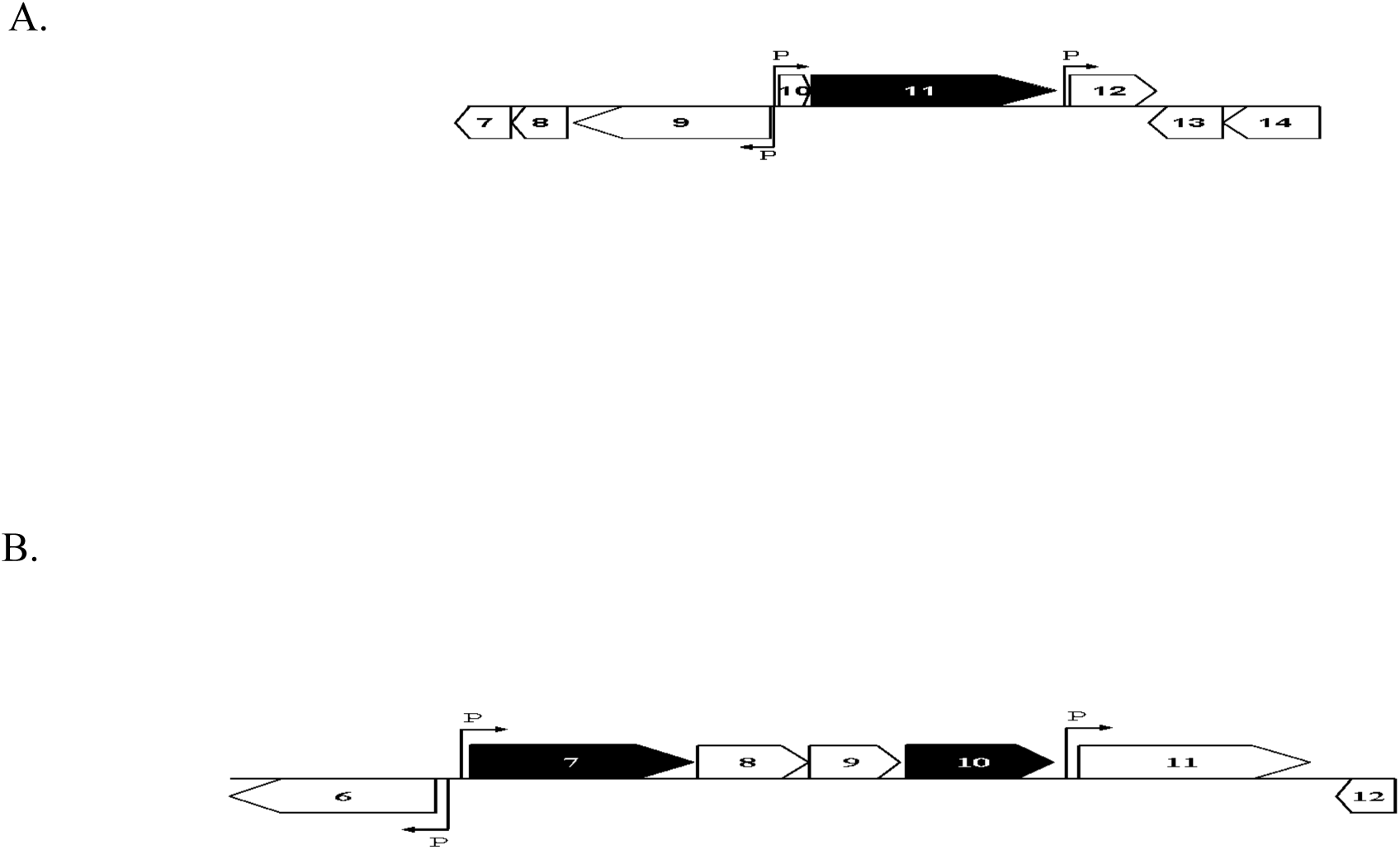
A DNA fragment carrying genes encoding β-galactosidases in Lac cosmids Lac36W and Lac161. (A) A gene locus from cosmid Lac36W (GenBank, KF255993). Lac36W_07, cytosine/adenosine deaminase; Lac36W_08, hypothetical protein; Lac36W_09, glutaminyl-tRNA synthetase; Lac36W_10, hypothetical protein; **Lac36W_11, β-galactosidase**; Lac36W_12, methionine-S-sulfoxide reductase; Lac36W_13, hypothetical protein; Lac36W_14, LysR family transcriptional regulator. The locations of potential promoter regions (P) were showed. (B) A gene locus from cosmid Lac161 (GenBank, KF255994). Lac161_06, histidine kinase; **Lac161_07, β-galactosidase**; Lac161_08, hypothetical protein; Lac161_09, hypothetical protein; **Lac161_10, β-galactosidase**; Lac161_11, hypothetical protein; Lac161_12, host specificity protein. The positions of potential promoter regions (P) were showed.

